# Colistin resistance in *Acinetobacter baumannii* is driven by multiple genomic traits: Evaluating the role of IS*Aba1*-driven *eptA* overexpression among Indian isolates

**DOI:** 10.1101/2021.01.07.425695

**Authors:** Saranya Vijayakumar, Jobin John Jacob, Karthick Vasudevan, Baby Abirami Shankar, Maria Lincy Francis, Agilandeeswari Kirubananthan, Shalini Anandan, Karthik Gunasekaran, Kamini Walia, Indranil Biswas, Keith S Kaye, Balaji Veeraraghavan

**Affiliations:** Department of Clinical Microbiology, Christian Medical College, Vellore – 632004, India; Department of Medicine, Christian Medical College, Vellore – 632004, India; Division of Epidemiology and Communicable Diseases, Indian Council for Medical Research, New Delhi – 110029, India; Department of Microbiology, Molecular Genetics and Immunology, University of Kansas Medical Center, United States of America; Division of Infectious Diseases, University of Michigan Medical School

**Author notes:** Corresponding author: Dr. V. Balaji, Professor, Department of Clinical Microbiology, Christian Medical College, Vellore – 632 004, Tamil Nadu, India.

## Abstract

Colistin resistance in *Acinetobacter baumannii* is mediated by multiple mechanisms. Recently, mutations within *pmrAB* two component system and overexpression of *eptA* due to upstream insertion of IS*Aba1* play a major role.To characterize colistin resistance mechanisms among the clinical isolates of *A. baumannii* in India. A total of 224 clinical isolates of *A. baumannii* collected from 2016 to 2019 were included in this study. Mutations within lipid A biosynthesis and *pmrAB* genes were characterized by Whole Genome Shotgun sequencing. Twenty eight complete genomes were further characterized for insertional inactivation of *lpx* genes and the association of IS*Aba1*-*eptA* using hybrid assembly approach. Non-sysnonymous mutations like M12I in *pmrA*, A138T and A444V in *pmrB* and E117K in *lpxD* were identified. Four of the five colistin resistant *A*.*baumannii* isolates had insertion of IS*Aba1* upstream *eptA*. No *mcr* genes were identified.Overall, the present study highlights the diversity of colistin resistance mechanisms in *A. baumannii*. IS*Aba1*-driven *eptA* overexpression could be responsible for colistin resistance among Indian isolates of colistin resistant *A. baumannii*.

## Introduction

*Acinetobacter baumannii* is a major nosocomial pathogen which is responsible for wide range of infections and has been reported as a public health problem globally [1, 2]. Multi-drug and extensively-drug resistant *A. baumannii* have been reported world-wide which results in the paucity of treatment options against *Acinetobacter* infections [3]. Carbapenems are considered to be the last-line antibiotics for treating *A. baumannii* infections and more than 80% resistance to carbapenem was reported [4]. More than 60% mortality rates have been reported for the most common carbapenem resistant *A. baumannii* (CRAB) infections like blood stream infections (BSI) and hospital-acquired pneumonia (HAP) [5].

Recently, World Health Organization (WHO) categorized CRAB as “Priority one” pathogen in the global list of antibiotic-resistant bacteria for the development of new antibiotics [3]. The most common antimicrobials considered for treating CRAB include colistin-based, tigecycline-based and sulbactam-based combinations [6]. Unfortunately, increased usage of colistin for treating *Acinetobacter* infection results in resistance to these last-line drugs [7].

In *A. baumannii* colistin resistance is mediated by multiple mechanisms [8]. This includes, (i) Loss of lipopolysaccharide (LPS) production, due to mutations in lipid A biosynthesis genes like *lpxA, lpxC* and *lpxD*. In addition, insertional inactivation of *lpxACD* genes due to insertion element, ISA*ba11* causes both loss of LPS and increased colistin resistance [9, 10]. (ii) Point mutations in *pmrA* and *pmrB* genes of *pmrAB* two component system (TCS) results in decreased membrane permeability leading to colistin resistance [11]. (iii) Recently, Gerson et al reported the presence of phosphoethanolamine (pEtN) transferase *eptA*, a homologue of *pmrC* and insertion of IS*Aba1* upstream *eptA* results in overexpression and high colistin resistance [3] and (iv) Colistin resistance due to plasmid mediated pEtN transferase, *mcr* genes [12].

In this study, a total of 224 clinical isolates of *A. baumannii* were characterized for their susceptibility profile and multiple resistance mechanism that contributes to colistin resistance. In addition, subset of complete genomes were characterized to investigate the inactivation of *lpxA* or *lpxC* by IS*Aba11* and to decipher the presence of IS*Aba1* upstream *eptA*.

## Materials and methods

### Bacterial isolates

A total of 1214 consecutive non-duplicate clinical isolates of *A. baumannii* collected during January 2016 to December 2019 from the routine cultures of clinical samples, blood (n=314) and endo-tracheal aspirate (ETA) (n=900) were included in this study. All the isolates were identified up to the species level as *Acinetobacter baumannii calcoaceticus* complex (*Acb* complex) using conventional biochemical methods [13]. MALDI-TOF was used to confirm at the species level as *Acinetobacter baumannii*. Further confirmation of *Acb* complex as *A. baumannii* was performed targeting chromosomally encoded *bla*_OXA-51 like_ gene by PCR [14].

### Anti-microbial susceptibility testing (AST)

AST was performed for all the isolates against different classes of antibiotics by Kirby Bauer disc diffusion (DD) method and interpreted according to Clinical Laboratory Standards Institute (CLSI) guidelines [15]. The antibiotics tested includes ceftazidime, cefepime, piperacillin/tazobactam, cefoperazone/Sulbactam, imipenem, meropenem, aztreonam, amikacin, netilmycin, tobramycin, levofloxacin, tetracycline, minocycline, tigecycline and trimethoprim-sulfamethoxazole.

### Minimum inhibitory concentration (MIC) by broth micro dilution (BMD)

As recommended by CLSI 2017 guidelines, colistin MIC was determined for all the blood and ETA isolates using broth micro dilution (BMD) and interpreted accordingly [16]. *Escherichia coli* ATCC 25922 and *Pseudomonas aeruginosa* ATCC 27853 were used as quality control (QC) strains. Also, *mcr-1* positive *E. coli* was used as an internal control. Two in-house quality controls, *Klebsiella pneumoniae* BA38416 with MIC of 0.5 µg/ml and *Klebsiella pneumoniae* BA25425 with 16 µg/ml were also included in every batch of testing.

### Whole genome sequencing (WGS), assembly and annotation

A subset of 224 clinical isolates of *A. baumannii* (Blood = 117 & ETA = 107) were selected based on the susceptibility profile for further characterization by WGS. Genomic DNA was extracted using QIAamp DNA Mini Kit (QIAGEN, Hilden, Germany) according to the manufacturer’s instructions and WGS was performed. In brief, short read sequencing was performed using IonTorrent™Personal Genome Machine™(PGM) (Life Technologies, Carlsbad, CA) with 400-bp read chemistry or Illumina MiSeq as per the manufacturer’s instructions. Long read sequencing was performed using SQK-LSK108 Kit R9 version (Oxford Nanopore Technologies, Oxford, UK) using 1D sequencing method according to the manufacturer’s protocol. To obtain complete genome, hybrid assembly was performed for a subset of 28 genomes (Blood = 21 and ETA= 7) using long reads from MinION and short reads from either IonTorrent or Illumina as described previously [17].

All the genomes were assembled and annotated using the NCBI Prokaryotic Genome Annotation Pipeline (PGAP). Furthermore, downstream analysis was done using tools from Center for Genomic Epidemiology (CGE) server (http://www.genomicepidemiology.org/). Antimicrobial resistance genes (ARG) were using ResFinder 3.0 database (https://cge.cbs.dtu.dk/services/ResFinder/) [18]. Sequence type of the isolates was assigned by MLST 2.0 tool using the Oxford scheme (*gltA, gyrB, gdhB, recA, cpn60, gpi*, and *rpoD* genes) (https://cge.cbs.dtu.dk//services/MLST/) [19]. SNP based phylogenetic tree was constructed and meta-data like colistin susceptibility, chromosomal mutation profile and International Clones were added using iTOL (https://itol.embl.de/) [20].

### Mutation analysis of *lpxACD* and *pmrAB* genes

*In-silico* mutation analysis of genes involved in lipid A biosysnthesis pathway (*lpxA, lpxC* and *lpxD*) and *pmrAB* TCS (*pmrA* and *pmrB*) were determined in all the genomes using Blast analysis (https://blast.ncbi.nlm.nih.gov) and compared with the reference strain of *A. baumannii* ATCC 17978 (GenBank Accession Number CP000521). Other reference strains like *A. baumannii* AYE (GenBank Accession Number NC010410), *A. baumannii* ACICU (GenBank Accession Number NC010611) and ATCC 19606 (GenBank Accession Number CP046654) were included in the analysis to identify the genetic polymorphisms. Plasmid mediated colistin resistance determinant gene (*mcr)* were detected using Resfinder and by *in-silico* Blast analysis against the reference sequences of *mcr* genes reported so far [21]. In addition to the mutation analysis, all the 28 complete genomes were characterized manually for other colistin resistance mechanisms like insertional inactivation of *lpxA, lpxC* and *lpxD* genes due to insertion sequence, IS*Aba11* and over-expression of *pmrC* homologue, *eptA* due to upstream presence of IS*Aba1*.

### Nucleotide accession numbers

The Whole Genome Shotgun project has been deposited at DDBJ/ENA/GenBank under BioProject numbers PRJNA603876, PRJNA604897, PRJNA610496 and PRJNA610503. The complete genome project has been deposited at DDBJ/ENA/GenBank under accession numbers AB01 (CP040080), AB02 (CP035672), AB03 (CP050388), AB04 (CP040040), AB05 (CP040047), AB06 (CP040050), AB07 (CP035930), AB08 (CP038500), AB09 (CP038644), AB010 (CP040053), AB011 (CP040056), AB012 (CP040084), AB013 (CP040087), AB014 (CP050421), AB015 (CP050403), AB016 (CP040259), AB017 (CP050385), AB018 (CP050523), AB019 (CP050400), AB020 (CP050390), AB021 (CP050410), AB022 (CP050412), AB023 (CP050415), AB024 (CP050425), AB025 (CP050432), AB026 (CP051474), AB027 (CP050526) and AB028 (CP050401).

## Results

### Bacterial isolates and AST

All the study isolates (n=1214) were first identified as *Acb* complex and confirmed as *A. baumannii* by MALDI-TOF. *bla*_OXA-51_ like PCR re-confirmed *Acb* complex as *A. baumannii*. AST revealed that 99% (Blood, n=313 and ETA, n=899) of the study isolates were extensively drug resistant (XDR) and showed resistance to all the tested antibiotics. Of the remaining two isolates, one from ETA was pan-susceptible (AB01-Accession No. CP040080) while the other from blood was multi-drug resistant (MDR). The MDR (AB025 - Accession No.CP050432) isolate was resistant to cephalosporins, fluoroquinolones, aminoglycosides, tetracyclines, tigecycline and trimethoprim-sulfamethoxazole while susceptible to carbapenems.

### Determination of colistin MIC by BMD

Among the tested isolates, 10.8% (n=34) from blood and 8.2% (n=74) from ETA were colistin resistant *A. baumannii* respectively. The colistin MIC range, MIC_50_ and MIC_90_ of the blood and ETA isolates were tabulated (Table 1).

**Table 1.** MIC range, MlC_50_ and MJC_90_ of colistin for clinical isolates of *A. baumannii*

| **Colistin MIC (µg/ml)** | **Blood (n=314)** | **Respiratory (n=900)** |
| --- | --- | --- |
| MIC range | 0.12 - 64 | 0.12 - 64 |
| MIC<sub>50</sub> | 0.5 | 1 |
| MIC<sub>90</sub> | 2 | 2 |

### Characterization of various colistin resistance mechanisms using WGS

A subset of 224 clinical isolates of *A. baumannii* was subjected to WGS based on the susceptibility profile. Of the 117 blood isolates, one isolate was MDR, 84 were carbapenem resistant-colistin susceptible *A. baumannii* (CR-ColSAB) and 32 were carbapenem -resistant-colistin resistant *A. baumannii* (CR-ColRAB). Similarly, among the 107 ETA, one was pan-susceptible, 46 were CR-ColSAB while 61 were CR-ColRAB.

Sequencing of *lpx* genes in both CR-ColSAB (n=131) and CR-ColRAB (n=93) revealed the presence of various amino acid substitutions. Within *lpxA*: Y131H, *lpxC*: C120R-P148S-F230Y-N287D and *lpxD*: Q4K-V63I-V93I-E117K-G166S-T287I-S299P substitutions were identified in comparison with the reference strain ATCC 17978. Three non-synonymous mutations Q4K, V63I and E117K within *lpxD* were identified in both CR-ColSAB and CR-ColRAB. Y131H, C120R and N287D substitutions were identified in all the three reference strains and all the CR-ColSAB and CR-ColRAB. Other amino-acid substitutions like P148S and F230Y in *lpxC*, V93I, G166S, T287I and S299P in *lpxD* were first described in this study. On the other hand, sequencing of *pmrAB* genes identified the following amino acid substitutions, *pmrA*: M12I and *pmrB*: A138T-L168S-A226T-V300E-G315A-P360Q-A444V. Single non-synonymous mutation, M12I in *pmrA* and A138T in *pmrB* was identified in all the CR-ColSAB and CR-ColRAB (Fig 1). None of the study isolates harbor any of the reported plasmid mediated colistin resistance determinant, *mcr*-1 or its homologues.

**Fig 1.**
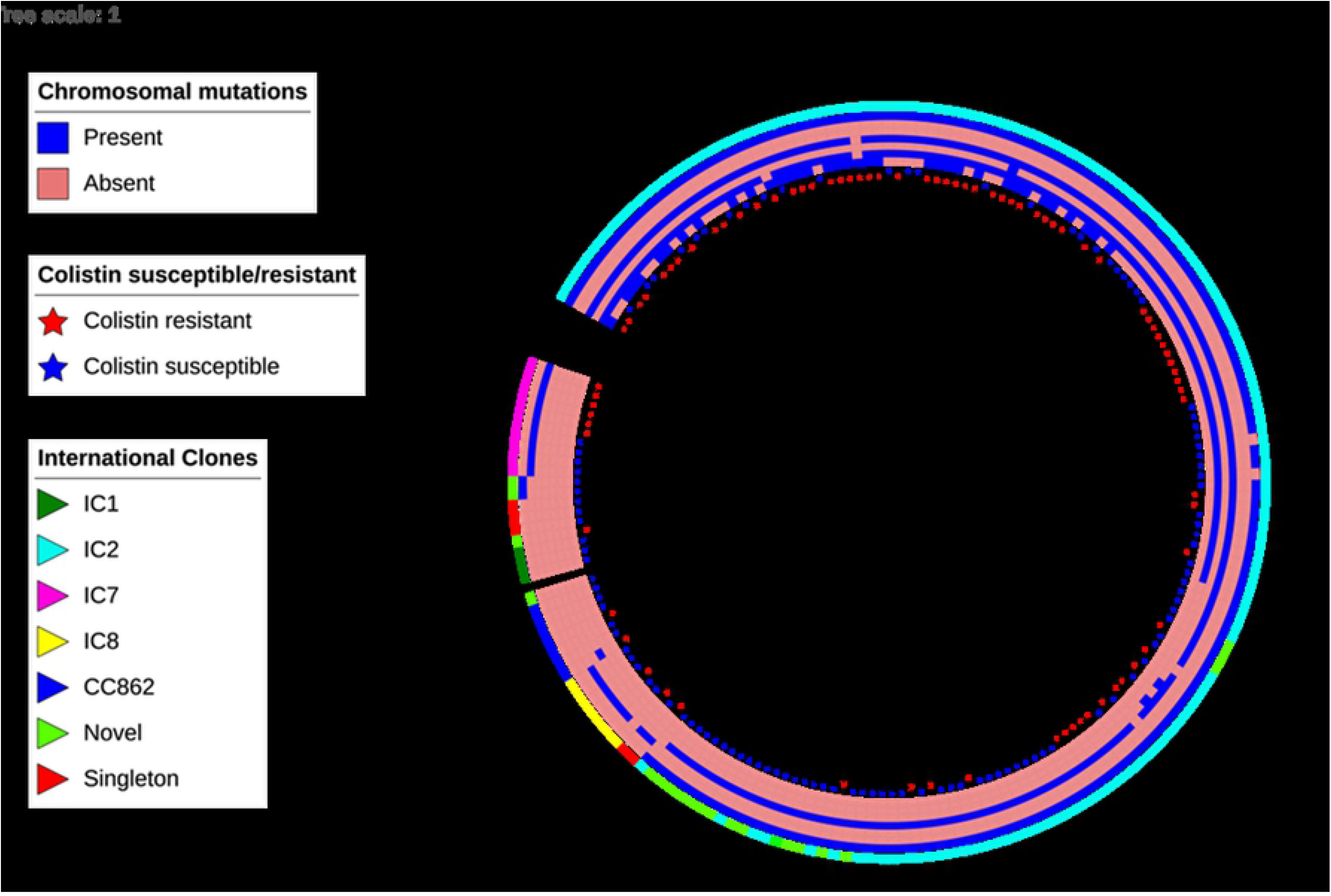
Phylogenetic tree of clinical isolates of colistin resistant and colistin susceptible *A. baumannii*. The outer ring shows isolates belonging to different International Clones (ICs), the middle rings depicts chromosomal mutations in both colistin resistant and susceptible isolates and the stars in the inner ring represents the colistin susceptibility of the clinical isolates of *A. baumannii*

Among the 28 complete genomes of *A. baumannii*, 21were from blood of which 17 were CR-ColSAB while four were CR-ColRAB. Of the seven from ETA, five were CR-ColSAB, one was CR-ColRAB and one was pan-susceptible. Characterization of complete genomes for other colistin resistance mechanisms showed the presence of IS*Aba11* among two CR-ColSAB blood isolates (AB016-Accession No. CP040259 & AB017-Accession No. CP050385), but no disruption of *lpxACD* observed. Atleast one copy of *eptA* was identified among 10 and 6 genomes of CR-ColSAB from blood and ETA respectively whereas more than one copy was present in 12 genomes (7 CR-ColSAB and 4 CR-ColRAB from blood while one CR-ColRAB from ETA). IS*Aba1* was found in opposite orientation upstream *eptA* in two genomes of CR-ColRAB from blood (AB02-Accession No. CP035672 & AB05-Accession No. CP040047) and in one CR-ColRAB from ETA (AB03-Accession No. CP050388). IS*Aba1* was present in direct orientation upstream *eptA* in one CR-ColRAB blood isolate (AB04-Accession No. CP040040). Two CR-ColSAB from blood have IS4 transposase but not upstream *eptA* (AB010-Accession No. CP040053 & AB011-Accession No. CP040056) whereas four CR-ColSAB from blood (AB08-Accession No. CP038500, AB09-Accession No. CP038644, AB025-Accession No. CP050432 and AB015-Accession No. CP050403) have other family transposases (Fig 2 − 3). In one CR-ColRAB (AB06-Accession No.CP040050), only amino-acid substitutions, Y131H in *lpxA*, C120R and N287D in *lpxC*, P148S and G315D in *lpxD* were identified while the other colistin resistance mechanisms were absent. The overall colistin resistance mechanisms characterized among the 28 hybrid genomes were described in (Table 2.)

**Table 2.**
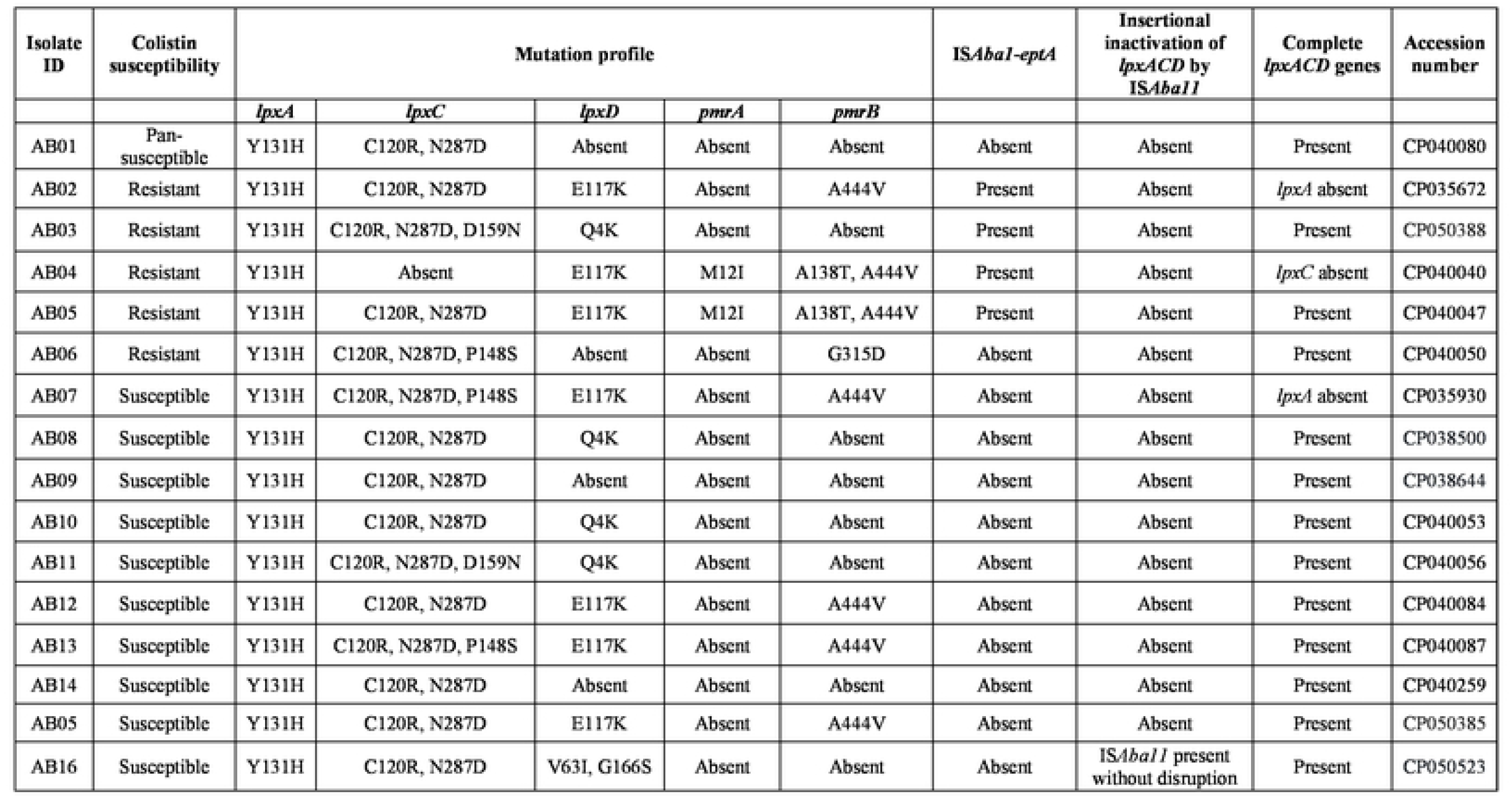
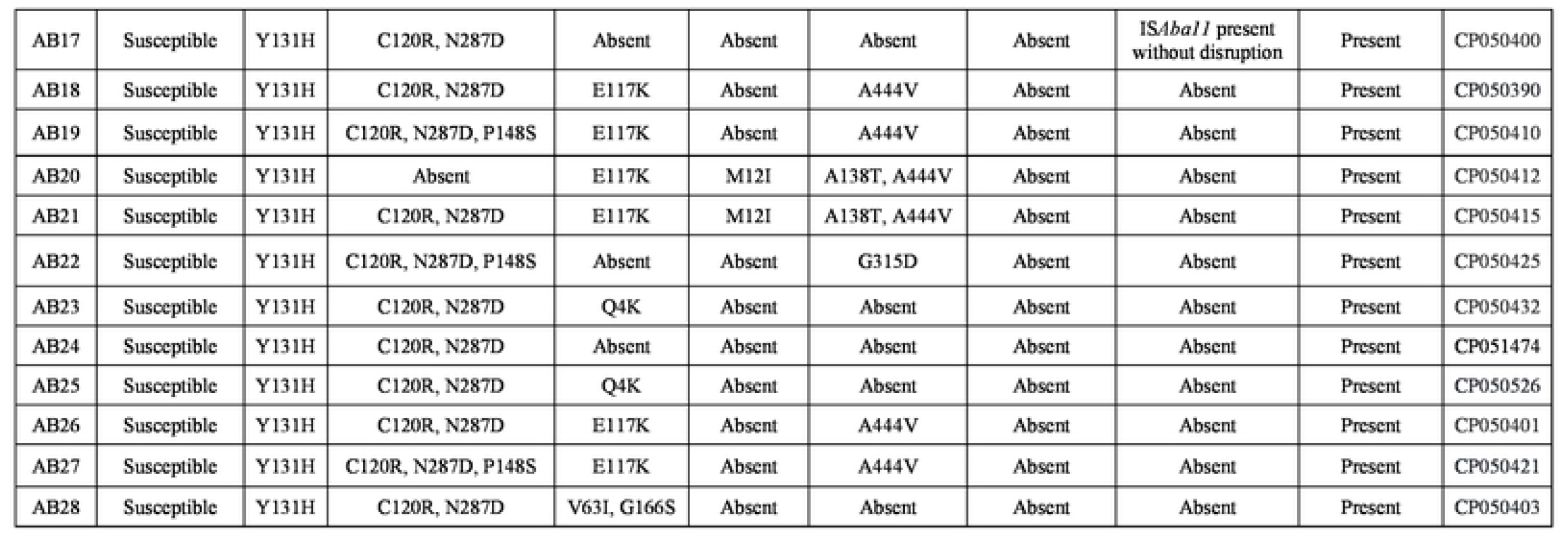
Cumulative findings of various colistin resistance mechanisms among complete genomes of *A. baumannii* (n=28)

**Fig 2:**
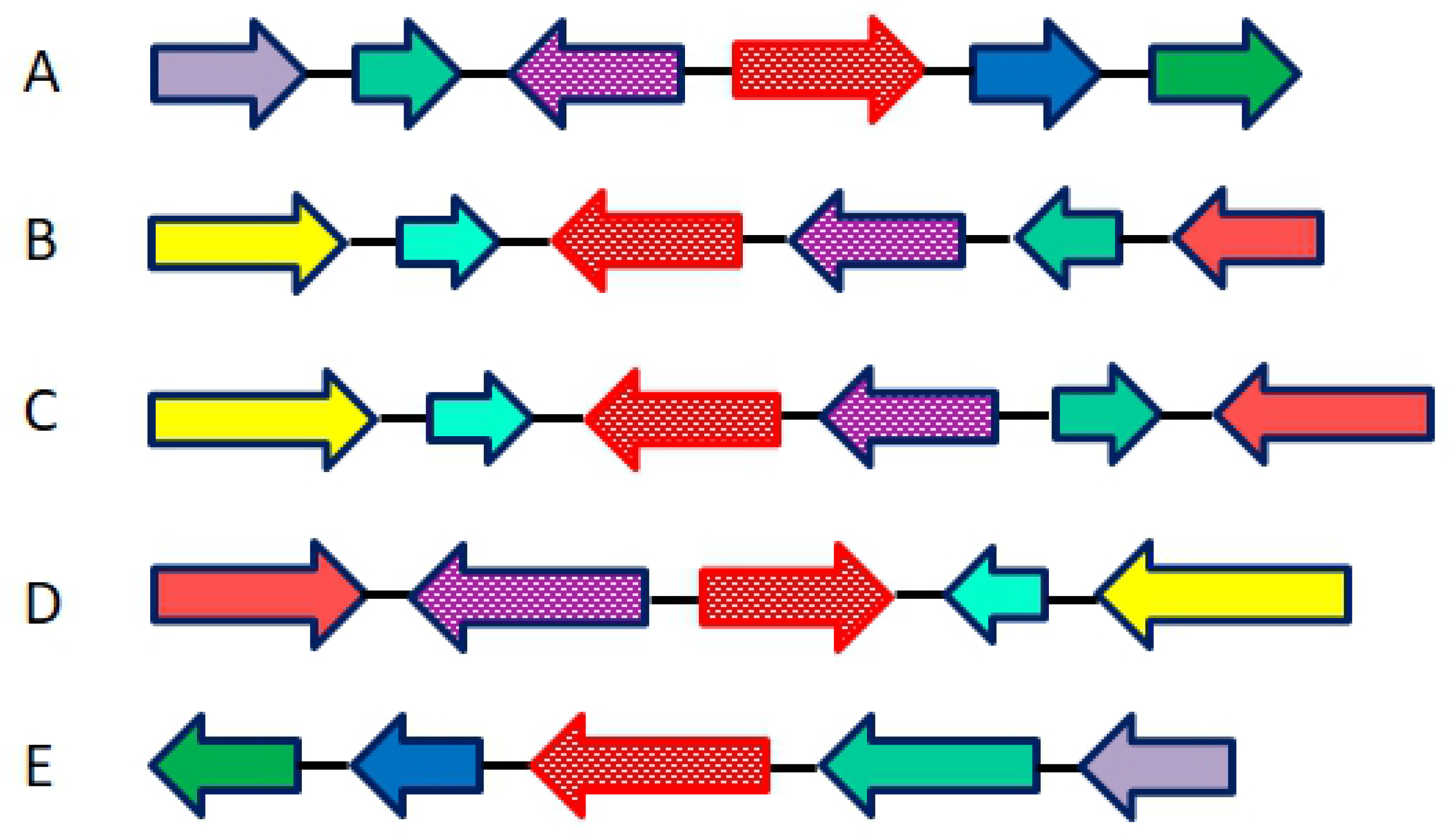
Graphical representation of genetic arrangement of *pmrC* homologue, *eptA* with upstream presence of IS*Aba1* among complete genomes of colistin resistant *A. baumannii* (A, B, C and D). The direction of arrow represents the orientation, *eptA* are shown as red-dotted arrow, IS*Aba1* as purple-dotted arrow, *pmrA* as blue arrow, *pmrB* as green arrow. The genetic arrangement of isolate E has *eptA* without IS*Aba1*

**Fig 3:**
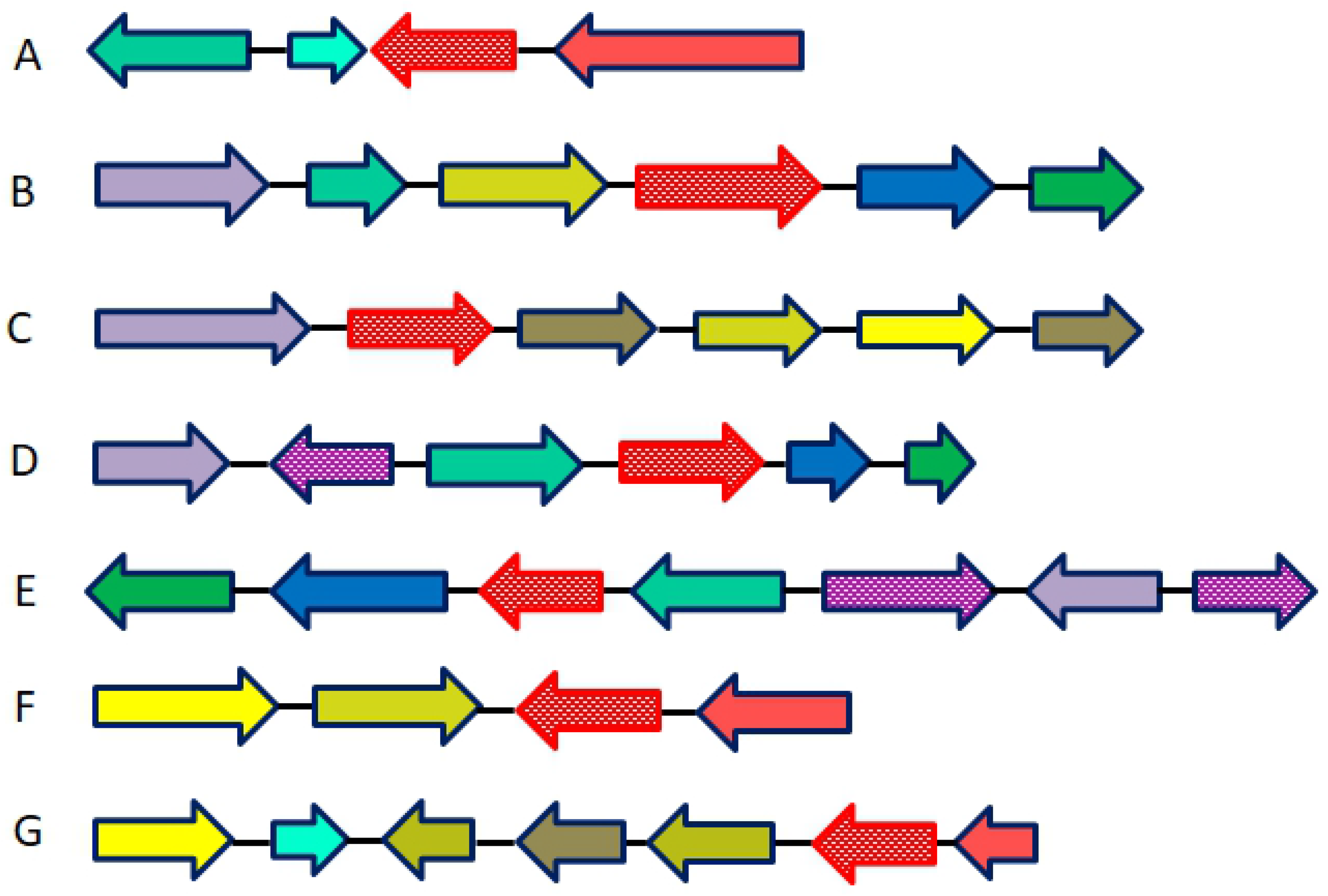
Graphical representation of genetic arrangement of *pmrC* homologue, *eptA* among complete genomes of colistin susceptible *A. baumannii*. The genetic arrangement of: A. *eptA* without insertion element, B. *eptA* with IS26 transposase, C. *eptA* with IS6 like element, D & E. *eptA* with IS4 transposase, F. *eptA* with IS256 transposase and G. *eptA* with IS66 transposase. The direction of arrow represents the orientation. *eptA* are shown by red-dotted arrow, IS26, IS6, IS256 and IS66 transposases as light green arrow, *pmrA* as blue arrows and *pmrB* as green arrow

### MLST

MLST Finder revealed that both CR-ColSAB and CR-ColRAB were classified into International Clones (ICs) - IC1, IC2, IC7, IC8, CC862 and singletons. Majority of the CR-ColSAB isolates (n=90) (69%) belongs to IC2 with diverse sequence types like ST848, ST349, ST451, ST218 and ST195. Whereas IC2 was the only lineage with ST848 among the CR-ColRAB isolates (n=73) (78 %). Lineage specific non-synonymous mutations like Q4K, V63I and E117K within *lpxD* belongs to IC8, IC7 and IC2 respectively. Single non-synonymous mutation, M12I in *pmrA*, A138T in *pmrB* and the combination of A138T-A444V within *pmrB* belongs to IC2.

## Discussion

Colistin is considered as one of the few therapeutic options available against CRAB infections and due to increased usage, colistin resistance rate is gradually increasing globally and becoming a healthcare concern [22]. In the current study, 8% to 11% colistin resistance has been observed. Reports from other countries showed increased colistin resistance rates of 16.7% from Bulgaria, 19.1% from Spain and 27% from Greece [23 – 25]. Therefore, it is important to explore and understand the mechanisms that contribute to colistin resistance which is an utmost need for routine surveillance [3]. Several mutations and genetic polymorphisms have been reported within *lpxACD* and *pmrAB* genes by various studies using WGS which is in concurrence with the current study [9, 10].

In this study, clinical isolates of CR-ColRAB and CR-ColSAB were characterized for multiple colistin resistance mechanisms. Mutation analysis in *lpxACD* identified three non-synonymous mutations; Q4K, V63I and E117K within *lpxD* in both CR-ColRAB and CR-ColSAB which concurs with previous studies [26, 27]. Recent studies report that mutations in *pmrB* as the major contributor for colistin resistance in *A. baumannii* [24, 26, 28, 29].The current study identified a non-synonymous mutation, A138T in *pmrB* while previous studies report other amino-acid substitutions together with A138T which is contrary to the current study [24, 26, 29]. In *pmrA*, a single non-synonymous mutation, M12I was first identified by Arroyo et al who also reported the association of M12I with colistin hetero-resistance [24, 26, 30]. In contrast, other studies reported G54E substitution within *pmrA* which was not identified in this study [29, 31]. Several studies reported the presence of mutations in *lpxD* along with *pmrB* genes [29]. In this study, both CR-ColRAB and CR-ColSAB has *lpxD* mutation, E117K in combination with A138T-A444V mutation in *pmrB*. This observation indicates that mutations in *lpxD* alone and *pmrB* alone may not be sufficient to induce colistin resistance and support the presence of synergistic activity of mutations within these genes in promoting colistin resistance [26].

Two MLST schemes, PubMLST (Oxford) and Pasteur MLST are available for *A. baumannii* [32]. Earlier studies reported IC2/global clone 2 (GC2) as the predominant lineage associated with outbreaks [8]. This finding correlates with the current study and identified in both CR-ColRAB and CR-ColSAB In the present study, IC1/GC1was identified among CR-ColSAB which is contrast to Snyman et al where GC1 was identified in both CR-ColSAB and CR-ColRAB [33]. Interestingly, lineage specific non-synonymous mutations, such as Q4K which belongs to IC8, V63I which belongs to to IC7 and E117K-M12I-A138T which belongs to IC2 were observed in this study and such findings have not been previously reported.

Moffatt et al was the first to report on the role of IS*Aba11* which causes insertional inactivation of *lpxA* or *lpxC* and leads to loss of LPS production and colistin resistance [10]. Mutations in *lpxC* gene due to insertion of IS*Aba11* and increased colistin resistance were recently reported by Marta et al [34].The current study revealed the presence of IS*Aba11* in two CR-ColSAB without disruption of *lpxA* or *lpxC* which is inconsistent with the previous studies.

*eptA* is a *pmrC* homologue with pEtN transferase activity [8]. In this study, *eptA* was present in both CR-ColSAB and CR-ColRAB and comparable with previous studies [3, 7 – 8, 35]. However, the presence of IS*Aba1* upstream *eptA* was identified only among the CR-ColRAB which results in overexpression of PetN transferase encoded either by *pmrC* or *eptA* and responsible for colistin resistance. Such findings are comparable with other studies whereas another study observed in both CR-ColSAB and CR-ColRAB [3, 7 – 8]. Presence of other family transposases were noticed among the CR-ColSAB which could be novel and further studies are necessary. Other colistin resistance mechanisms like disruption of *eptA* by IS*Aba125*and increased expression of *eptA* due to the presence of IS*Aba1* in reverse orientation upstream *eptA* were reported recently [3].Though difference in the orientation of IS*Aba1* with respect to *eptA* was identified in this study, no expression studies were done to confirm the same.

In 2016, plasmid mediated PetN transferase, *mcr-1* was identified from China that contributed to colistin resistance to *E. coli* [36]. Currently, nine *mcr* types (*mcr* 1-9) and approximately 56 *mcr* variants are available in the GenBank database [37]. Though recent studies reported the presence of *mcr-4*.*3* in *A. baumannii* isolated from pig feces, food sample and clinical strains, none of the current study isolates harbor *mcr* genes [37 – 39].

Though colistin-based combinations can be considered for treating CRAB infections, various controversies have been reported recently with the usage of polymyxins [40].Two global organizations, CLSI and EUCAST revised the interpretive criteria for *in-vitro* polymyxin susceptibility testing and suggested to prefer non-polymyxin agents for treating *Acinetobacter* infections [41]. Such revision would effectively helpful in considering polymyxin as a treatment option in selected cases [41]. Further clinical trials to determine the efficacy of novel agents like cefiderocol and eravacycline against BSI and VAP are recommended [42]. None of the newly available drug combinations have clinical activity against CRAB infections except for the novel β-lactam-β-lactamase enhancer, cefepime-zidebactam [6, 42].

## Conclusion

The current study provides characterization of multiple resistance mechanisms that could be responsible for colistin resistance among clinical isolates of *A. baumannii*. Previously reported non-synonymous mutations as well as other amino-acid substitutions that are not described previously within *lpxD, pmrA* and *pmrB* genes were identified. Presence of *lpxD* mutation, E117K along with *pmrB* mutations A138T-A444V indicates the synergistic activity of mutations and results in colistin resistance. Recently, *mcr-4*.*3* on plasmids has been identified among clinical isolates of *A. baumannii* which is alarming. Two colistin susceptible isolates harbored IS*Aba11* and studies to understand the role of IS*Aba11* towards colistin resistance is essential. Though *pmrC* expression and addition of pEtN transferase to lipid A was regulated by *pmrAB* TCS, that alone doesn’t considered as a sole contributor of colistin resistance. The presence of *eptA* was quite common and insertion of IS*Aba1* upstream *eptA* among colistin resistant isolates could be associated with colistin resistance. The exact resistance mechanism that contributes to colistin was not elucidated for one isolate and requires further investigation. Overall, the present study highlights the diversity of colistin resistance mechanisms described in *A. baumannii* and for complete understanding further extensive studies are necessary.

## Acknowledgements

The authors gratefully acknowledge the Institutional Review Board of the Christian Medical College, Vellore (83-i/11/13) and Indian Council of Medical Research, New Delhi, India (ref. no AMR/TF/54/13ECDHIII dated 23/10/2013). The study is a part of the PhD. Dissertation under The Tamil Nadu Dr. M.G.R. Medical University.

## Transparency declarations

None to declare

